# Lateral Entorhinal Cortex Dysfunction in Alzheimer’s disease Mice

**DOI:** 10.1101/2024.04.15.589589

**Authors:** Radha Raghuraman, Andrew Aoun, Mathieu Herman, Oliver Shetler, Eden Nahmani, S. Abid Hussaini

## Abstract

In Alzheimer’s disease (AD), the formation of amyloid beta (A*β*) and neurofibrillary tangles (NFTs) leads to neuronal loss in entorhinal cortex (EC), a crucial brain region for memory and navigation. These pathological changes are concurrent with the onset of memory impairments in AD patients with symptoms of forgetfulness such as misplacing items, disorientation in familiar environments etc. The lateral EC (LEC) is associated with non-spatial memory processing including object recognition. Since some LEC neurons fire in response to objects (object cells) while others fire at locations previously occupied by objects (trace cells), pathology in this region could lead to dysfunction in object-location coding. In this paper we show that a transgenic AD mouse model, EC-App/Tau, which expresses both APP and tau primarily in the EC region, have deficits in LEC-specific memory tasks. Using in vivo single-unit electrophysiology recordings we show that LEC neurons are hyperactive with low information content and high sparsity compared to the controls indicating poor firing fidelity. We finally show that object cells and trace cells fire less precisely in the EC-App/Tau mice compared to controls indicating poor encoding of objects. Overall, we show that AD pathology causes erratic firing of LEC neurons and object coding defects leading to LEC-specific memory impairment.

## Introduction

Detecting early signs of neuronal vulnerability in Alzheimer’s disease (AD) prior to the onset of cognitive symptoms is crucial for interventions to prevent further progression. In AD, the entorhinal cortex (EC) is shown to have selective vulnerability due to the accumulation of amyloid beta (A*β*) and tau, especially in the lateral EC (LEC) [1–3]. Therefore, early AD symptoms such as misplacing objects, anosmia and forgetting events could be due to the distinct role of LEC in object-, odor- and episodic memories [4–11]. Anatomical studies have shown that while hippocampal place cells receive spatial information from the medial EC (MEC), the non-spatial information is mainly provided by the LEC [12–15]. LEC neurons show very little spatial selectivity in an open field, even with several cues and landmarks, but fire in response to discrete objects [13, 16–19]. Two distinct types of specialized cells viz., object cells and object-trace cells have been identified in the LEC of rats thus far [9]. While an object cell fires in response to an object, an object-trace cell fires at a location previously occupied by an object, indicating that there are two independent neural correlates for object-place memory in the LEC [9]. As part of non-spatial memory, these cells collect, remember and process object-place information to be integrated with spatial memory in the hippocampus.

The LEC also processes olfactory information and receives input from both olfactory bulb (OB) and piriform cortex (PCX). The LEC neurons, unlike the PCX neurons, are more narrowly tuned and process complex odors [20]. The dominant input to LEC is from the afferent fibers of olfactory areas [21] which then project back to both the aforementioned areas [22, 23]. Odor-discrimination-trained animals display EC signaling to OB prior to the odor onset which prepares the system for odor sampling [24]. This indicates LEC to be instrumental in providing highly odor-specific, expectation-dependent feedback to the olfactory bulb [25, 26]. Any loss or deterrence of this feedback results in the interference with odor memory, thereby contributing to early impairment in olfactory perception that is commonly observed in AD [27–30]. In addition, the LEC is known to coalesce multisensory information. Several studies, including LEC lesions, have shown that object-context or odor-context memory is impaired while the animal’s ability to discern new objects, odors and contexts individually is intact, indicating that LEC is critical in complex, association memory and plays an important role in the integration of episodic information [5, 6, 31–34].

Several studies including ours have shown that amyloid beta and tau pathologies affect MEC neurons and impair grid and speed cell functions, but very little is known about its impact on LEC neuronal function during behavior [35–38]. Functional imaging studies have shown that expressing App and tau in the EC of mice affects LEC function. Further, in vitro studies in App knockin mice have found LEC neurons to be vulnerable, with increased neuronal excitability compared to other brain regions [3, 8]. While we have previously described object and trace cells in mice ([39], symposium), to our knowledge, this is the first study that investigates object and trace cells in LEC of AD mice, and how AD pathology impacts them. In this paper we show that AD pathology in the LEC leads to neuronal hyperactivity with loss of firing fidelity and poor encoding of objects possibly leading to object-context memory deficits.

## Results

### EC-App/Tau Mice Show LEC-specific Memory Impairment

EC-App/Tau mice accumulate A*β* and tau in the entire hippocampal formation but a functional imaging study showed that the most vulnerable region was the LEC ([3]). In the current study, using the same mouse model we confirmed that A*β* and tau accumulate in the hippocampal formation with higher pathology in the EC—a result of higher A*β* and tau expression driven by the neuropsin promoter (Figure S1). Since this increased pathology in LEC could possibly result in LEC-specific memory deficits, we tested the memory of 20-month-old EC-App/Tau mice and age-matched Neuropsin control (NON) mice using three behavioral tasks. We used the Object context recognition (OCR) and Odor discrimination (OD) task to test LEC-specific memory, and the T-maze alternation task to test the hippocampal-dependent memory ([4, 5, 33, 40, 41]). In the OCR task, the mice explored two identical sets of objects (two cubes or two triangular pyramids) across two different contexts (black cylinder or gray striped cylinder) during the 2-day training phase. During the test phase on day 3, one of the objects in one context was replaced with an object belonging to the other context (e.g. a triangular pyramid replaced with a cube in the gray striped context). The expectation was that the mice with intact memory would spend more time exploring the out-of-context ‘novel’ object (blue cube) (Figure 1, Left). The EC-APP/tau mice did not explore the novel objects as much as the controls, evident from the Fisher’s exact test (*p* < 0.001), wherein the controls show a significant difference with the EC-App/Tau in being able to recognize the objects in the context. In the OD task, mice were trained to turn left or right to receive food pellets based on the odor they were presented with in the stem of the T-maze (Figure 1, Right). EC-APP/tau mice made many errors compared to the controls, indicating poor odor discrimination (evident from the contingency table representing the frequencies of corresponding arm choices made by the mice belonging to respective cohorts of EC-App/Tau: incorrect arm choice = 68.89% and correct arm choice = 31.11%) while the NON group shows 40% incorrect arm choice and 60% correct arm choice from Fig1 right panel representing odor discrimination task). Finally, we tested mice in a T-maze spontaneous alternation task where mice were trained to turn left and right by blocking one arm in the sample trial and allowing them to choose between the two arms in the choice trial. While the EC-App/Tau mice made more errors in this task with a greater amount of incorrect alternations compared to the control mice, the effect was not significant (p = 0.1178) (Figure S2).

**Figure 1.**
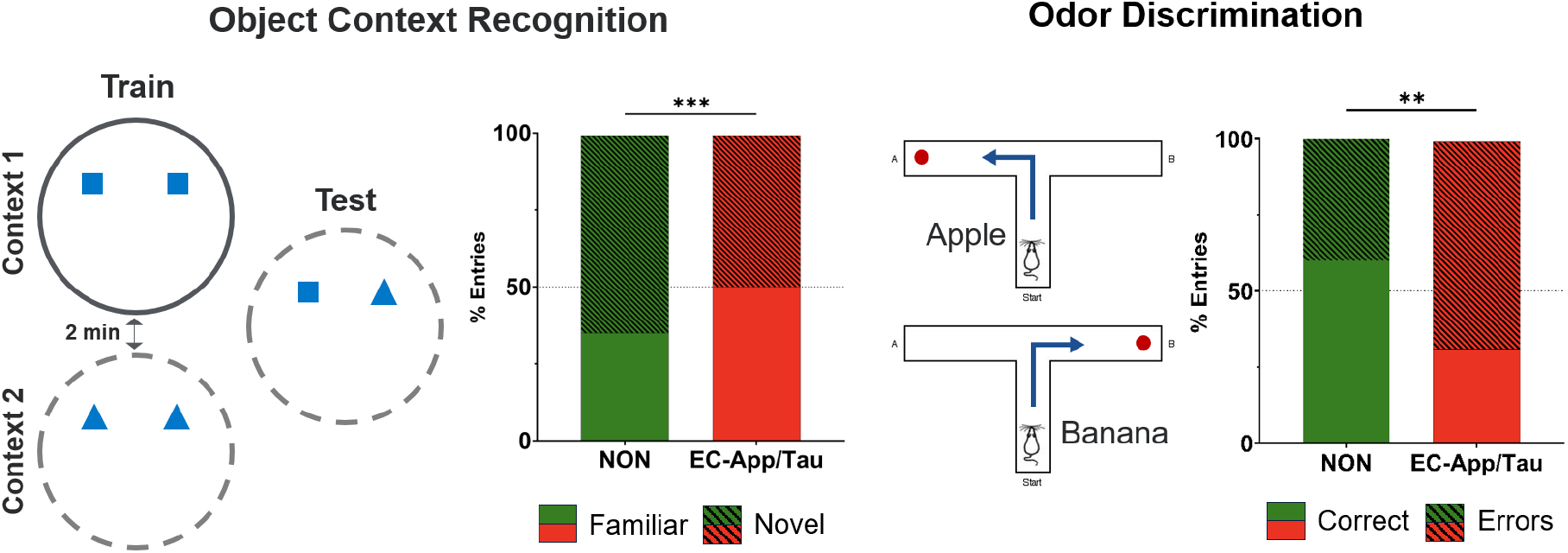
Object Context Recognition and Odor Discrimination tasks. The left panel shows the set up of Object Context Recognition (OCR) task with context 1 showing a plain cylindrical arena with two identical cube shaped objects and context 2 showing a striped cylindrical arena showing two identical pyramid shaped objects in the training phase. In the test phase, a cube and a pyramid object were presented to the mice in one of the two contexts. The adjacent histogram represents the frequencies (in percentage) indicating the entries made by the mice in each object zone. The right panel shows the set up of Odor Disrimination (OD) task where the mice are trained to turn left when presented to apple odor and turn right when presented with banana odor. A sugar pellet was provided as a reward on successful discrimination. The histogram on the right shows the contingency table representing the frequencies corresponding to the correct (in plain) and incorrect arm choices (represented in hatched lines) made by the mice from the respective cohort

### LEC Neurons Show Increased Firing and Reduced Spatial Information

Since LEC-specific memory in EC-App/Tau mice was affected, we focused on understanding how A*β* and tau impact the neurons in the LEC. We collected single-unit activity from the LEC neurons of: 22-month-old EC-App/Tau mice, age-matched NON mice, and 8-month-old C57BL/6J (B6) mice (Figure S7 tetrode location), while they explored an open field cylindrical arena with or without an object for 10 minutes (see next section and methods for details). We found that the LEC neurons of EC-App/Tau mice had a higher firing rate (12.79 Hz.) compared to the NON mice (1.81 Hz.) and B6 mice (0.93 Hz.) (Figure2, left), and NON mice had higher firing rate compared to B6 mice (Kruskal Wallis test: *p* < 0.0001 for all comparisons). The spatial information content in EC-App/Tau mice was lower compared to the control mice indicating poor encoding of spatial information (Figure 2, right). The information content in EC-App/Tau mice was 0.44 bits/spike compared to NON mice (1.38 bits/spike) and B6 mice (1.28 bits/spike) (Kruskal Wallis test: *p* < 0.0001). While it is noteworthy that there was no significant difference between the NON and B6 mice (Kruskal Wallis test: *p >* 0.99). Consistent with this, the sparsity of LEC neurons in EC-App/Tau mice was higher compared to the controls (Figure S3). However, both selectivity and coherence of EC-App/Tau mice were similar to B6 mice (Figure S3). These impaired firing properties of LEC neurons in EC-App/Tau mice indicated that their responses towards objects might be affected too.

**Figure 2.**
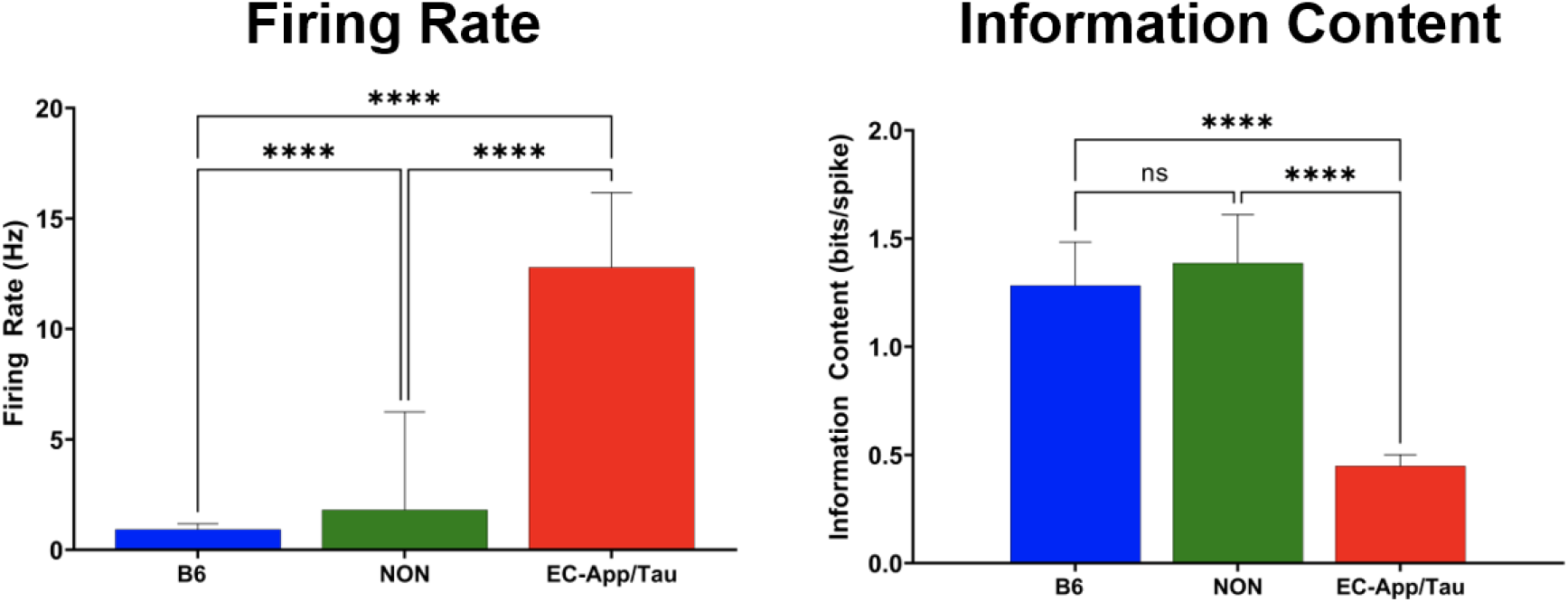
Firing Rate and Skaggs Information Content The left panel represents the histogram showing comparisons of firing rate amongst the three cohorts - B6 (blue), NON (green) and EC-App/Tau (red). The right panel represents the histogram showing comparisons of Skagg’s information content amongst the three cohorts - B6 (blue), NON (green), and EC-App/Tau (red). P values are represented by asterisks and error bars are SEM

### Object and Trace Cells in EC-App/Tau Mice Are Poorly Tuned

Studies in rats have shown that LEC neurons fire in response to objects (object cells) and at locations occupied by previous objects (object-trace or trace cells) ([9, 17]). We report object cells and trace cells for the first time in mice with varying degrees of tuning towards objects (black circles) and trace objects (white circles) due to aging and AD (3). Since the firing of EC-App/Tau LEC neurons were impaired (figure 2 and figure S2), their responses towards object locations could also be affected, explaining the object memory impairment in these mice (1). We recorded from LEC neurons as mice explored a cylindrical arena for 3 or more sessions for 10 minutes each with 1 minute inter-trial interval. In the first session, the mice explored the arena for 10 minutes without an object (3, Object cells, column 1) and in the subsequent two consecutive sessions, an object was positioned at the edge of the cylinder at one of the following locations relative to the center: 0 degrees (North), 90 degrees (East), 180 degrees (South) or 270 degrees (West) (3, Object cells, columns 2-3).

We quantified LEC neuronal responses towards objects in each session using the EMD (Earth Mover’s Distance) metric ([42]). The distances between firing fields and the object locations were computed using a single point EMD distance (see methods). For no-object sessions (NO), the center of the arena was used as a reference point. A cell whose firing field was closest to an object location compared to other tested object locations for 2 consecutive sessions was defined as an object cell. Representative firing fields for each group; B6, NON and EC-App/Tau, for no-object and objects at 4 possible locations (black circles in columns 2-5) are shown in figure 3, Object cells. A cell whose firing field was closest to a previous object location compared to other tested object locations was defined as a trace cell. Representative firing fields for each group for objects previously placed at 4 possible locations (white circles in columns 2-5) are shown in figure 3, Trace cells. We plotted the average population firing response of all neurons towards all 4 object locations combined (figure 3, Average) and individually (Figure S4) for each groups. The combined population response of LEC neurons revealed that the firing fields in B6 mice were more uniformly spread across 4 object locations, with high firing (warm colors) at the edge of the circle and low firing (cold colors) in the center, compared to NON and EC-Tau/App mice. Among the 3 groups, the average population response of LEC neurons in the EC-App/Tau mice was higher in the center, indicating poor tuning towards objects locations. The average population response also indicated that the quality of object and trace cells was vastly different across the groups.

**Figure 3.**
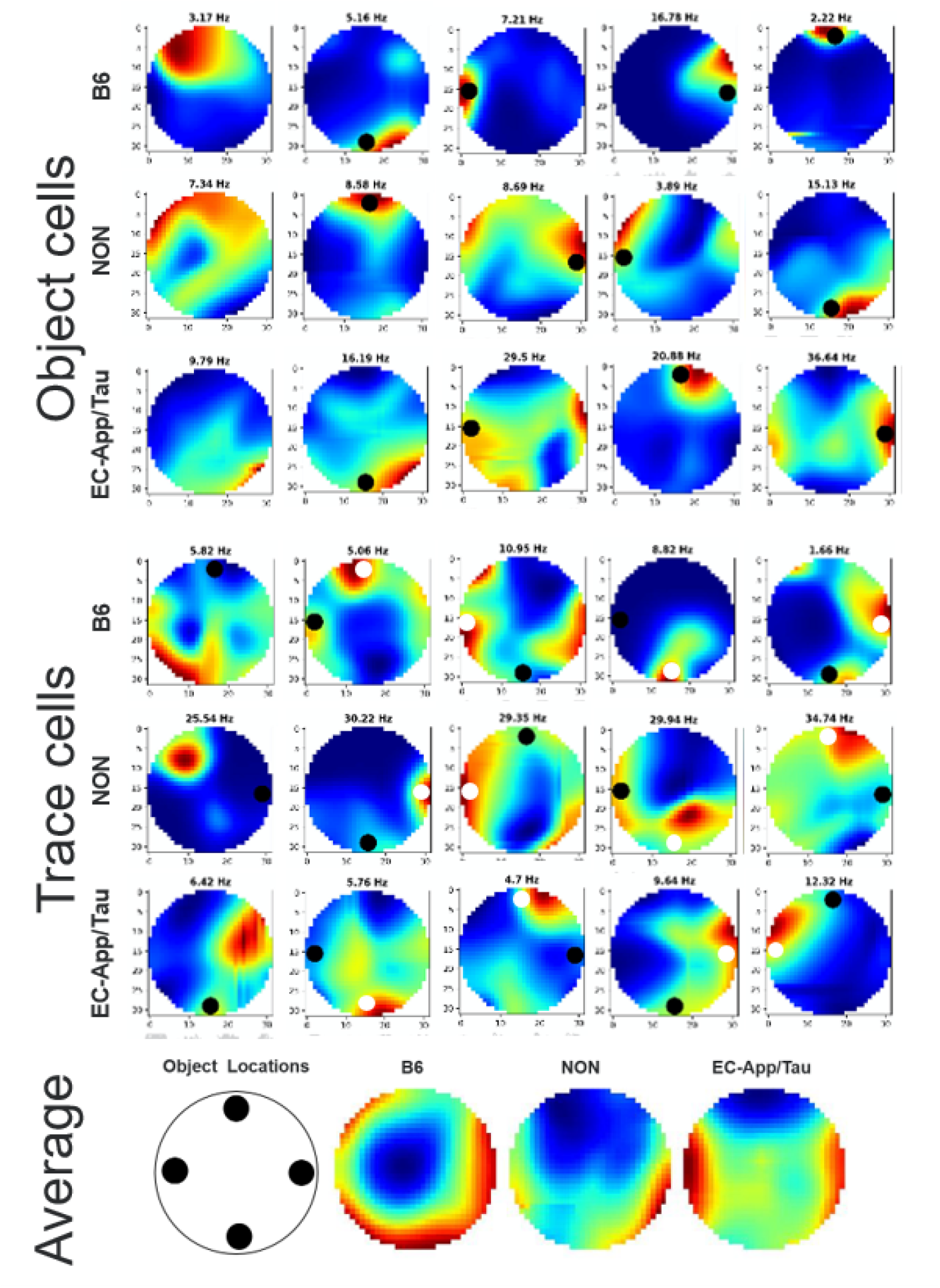
**Object Cells:** Representative object cell responses for B6, NON and EC-App/Tau mice for no object (left column) and 4 possible current object locations; 0, 90, 180 and 270 (black circles in columns 2-5). Columns 1-3 are part of the same session and rest from another session. **Trace Cells:** Representative trace cell responses for 4 possible previous object locations; 0, 90, 180 and 270 (while circles in columns 2-5). Columns 1 and 2 are part of the same session. Black circles indicate current object location. Average population response of neurons towards objects for the 3 groups.

### Object and Trace cells in EC-App/Tau Mice Are Less Stable

We measured the quantity and quality of object and trace cells in the 3 groups. We found about 5%-10% object cells (figure 4, top left), with significantly more cells in NON mice compared to the other two groups (B6: 5%, NON: 10% and EC-App/Tau:5%). There were about 20%-30% trace cells (figure 4, top right), with significantly more cells in NON compared to B6 mice (B6: 24%, NON: 30% and EC-App/Tau: 26%). We then quantified how many of the above cells are truly firing near the current or previous object’s location versus firing away from the object (figure 4, bottom), either closer to the center (bottom left) or in between two object positions (bottom right).

**Figure 4.**
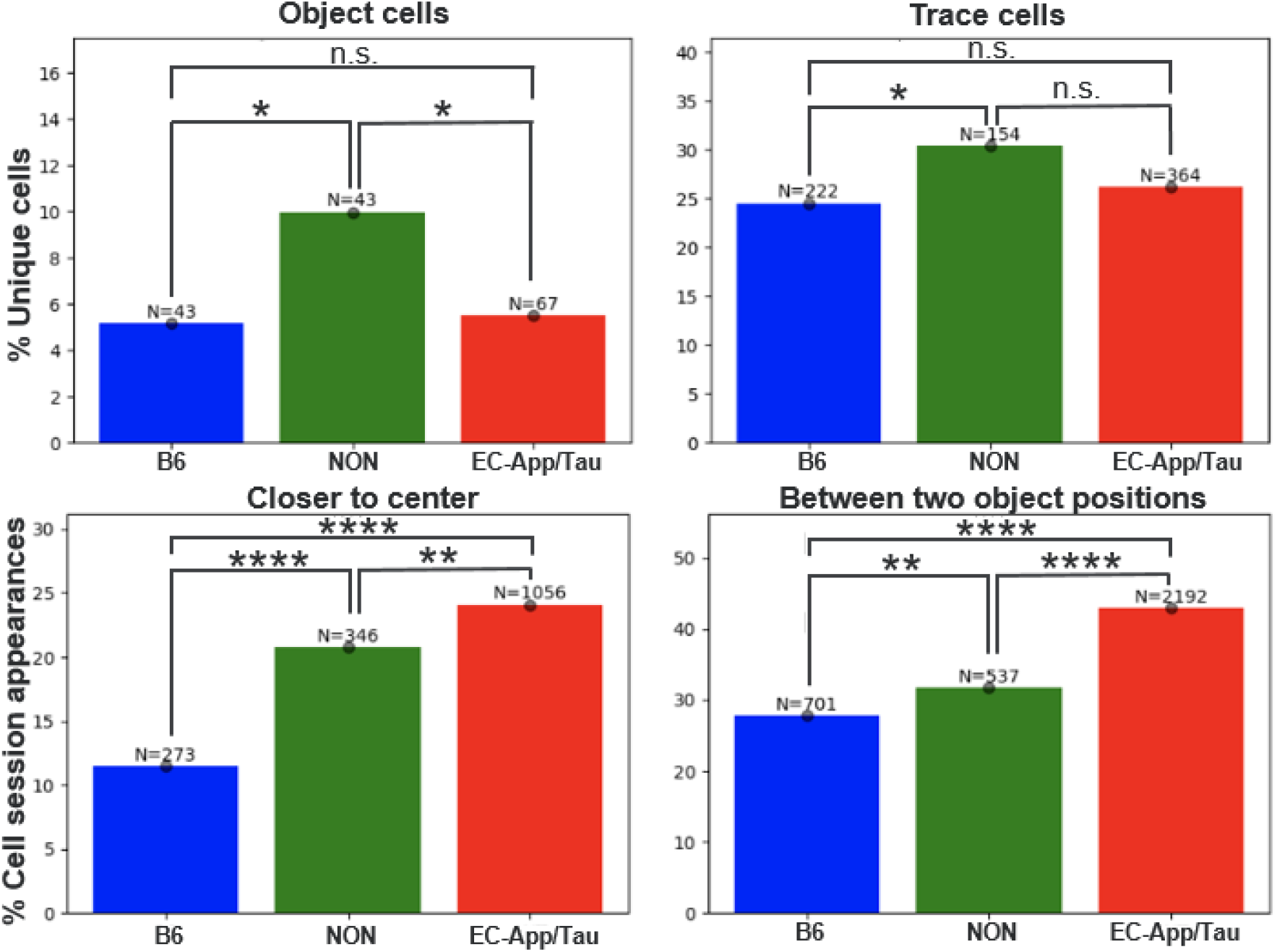
Cell Counts. Cell counts are compared using Fisher’s exact test. The percentage of cells which show object and trace behavior across their sessions compared for the 3 groups (top panel). The first quality metric shows the proportion of cases where the EMD quantile is smaller relative to the center point than to any of the 4 object locations (left). The second shows the proportion of cases between two object locations such that the reference point of the minimum quantile is different for the field-restricted EMD and the field centroid distance (right). Raw cell and cell-session counts are provided above each proportion.

We found that many object and trace cell sessions in the EC-App/Tau mice were of low quality, i.e. fields were farther away from current or previous object locations compared to NON and B6 mice. In about 23% of sessions, the fields of EC-App/Tau mice were closer to the center of the arena rather than any of the 4 corners compared to the fields of B6 (10%) and NON mice (20%). In about 40% of the sessions, the fields of EC-App/Tau mice were in between two object positions (for example at 45 degree position instead of 0 or 90 degrees) compared to the fields of B6 (25%) and NON mice (30%). Similarly, cell sessions in NON mice were of lower quality compared to B6 mice. The remaining 60%-75% of cells were not tuned to an object (unassigned cells).

To interpret group stability however, it was important to choose one reference point to remain consistent across groups. Given that we expected object and trace cell response, and that our tested locations were along the edges, we chose the center of the arena as our reference and hypothesized that greater object/trace content would shift stability away from the center. To compare stability, we fit regression models to the distribution of EMD quantiles associated with each group pair and for a given remapping metric. This model incorporated both random animal effects as well as fixed effects for group and session number. For both metrics, we observed the expected trend in stability with B6 showing the largest averaged quantile followed by NON and subsequently EC-App/Tau (Figure S5, top panel). We found that the fitted coefficient for the quantiles computed relative to the center of the arena indicated that the EC-App/Tau group was significantly less stable than B6 for the whole map EMD metric while both EC-App/Tau and NON were significantly less stable than B6 for the field-restricted EMD metric.

Importantly, unlike NON and EC-APP/Tau mice, we had few B6 recordings with more than 3 sessions, and therefore limited the net cell type counts across groups to 3 sessions (figure 6). However, given that LEC responses are shaped by learning and experience, we repeated the comparison between NON and the EC-APP/Tau mice for up to 6 consecutive sessions (Figure S5, bottom panel). This validates the use of our quality metrics as we found that the proportion of fields between two angles remained stable across sessions for both groups.

Additionally, we show that with more experience, NON mice increase their tuning by reducing the proportion of cells closer to the center while ANT mice increase it. To confirm this result, we show that the trend in EMD quantiles is significantly positive for the NON mice while that of the EC-APP/Tau is significantly negative or close to zero (Figure S5). We therefore repeated the object and trace counts after filtering out ambiguous cases based on our quality metrics. While this reduced the raw count of object and trace cells, EC-APP/Tau mice demonstrated a significantly reduced proportion of both cell types compared to the NON mice (Figure S6, top panel). This shows how learning and memory deficits in EC/App-Tau mice are specifically tied to the reduced quality of object and trace cell content.

## Discussion

This paper shows that AD pathology in EC-App/Tau mice impairs neuronal function in the LEC and leads to LEC-specific memory deficits. Previous study that characterized the EC-App/Tau mice ([3]), showed dysfunction in the LEC using fMRI but did not perform behavioral task. In this paper, among the three behavioral tasks, viz. Object context recognition (OCR), Odor discrimination (OD), and T-maze alternation tasks, the mice were affected only in the LEC-specific, OCR and OD tasks and not in the hippocampal-dependent task. This is consistent with previous studies that have shown OCR deficits due to lesions but intact novel object memory [5]. Our results also suggest that while there is amyloid beta and tau in other hippocampal regions, the EC is affected the most leading to LEC-specific memory impairment.

Previous studies have shown that A*β* and tau cause neuronal hyperexcitability leading to network dysfunction ([43–46]) and memory deficits. We confirm that the LEC neurons of EC-App/Tau mice have higher firing rate with low information content and high sparsity compared to the controls. This indicates poor firing fidelity, which could contribute to the observed memory deficits. The fMRI study ([3]) done on EC-App/Tau mice, which measured cerebral blood volume (CBV), showed hypofunction in the LEC region but we observe hyperexcitability in our data. This discrepancy could be due to the differences in recording method, age of mice and brain state. While CBV measures relative blood volume in the LEC our technique directly measures neuronal firing which might be inherently different. The age used in fMRI was 12 months while we used older, 22+ old mice. Finally, fMRI was done on anesthetized mice while our mice were freely moving. Indeed, a paper has shown that CBV and firing activity can be uncorrelated depending on resting state versus during tasks ([47]).

We found about 5-10% of object cells and 20-30% of trace cells from a total of 704 neurons.

Although we found an equal or greater number of object and trace cells in the EC-App/Tau mice compared to other groups, their coding for object location was poor (Figure 4 and S6 bottom panels). Using a previously described EMD metric ([42]), we show that the tuning and stability of object cells and trace cells in EC-App/Tau mice is significantly poorer compared to the controls. Many object and trace cell sessions in the EC-App/Tau mice were of low quality, with fields farther away from current or previous object locations. This poor tuning and stability might explain to the observed object-context memory deficits (Figure 1). Additional evidence for this is provided in the findings that, with experience, EC-App/Tau mice recruit fewer object and trace cells compared to controls (Figure S6, Top panel). Overall, our results using EMD metric reveal that the tuning for objects and the stability of tuning across sessions is significantly affected in EC-App/Tau mice compared to control mice.

One of the caveats of the EC-App/Tau model used here is the known leakiness in the expression of neuropsin which could express amyloid beta and tau in other regions too [48]. While we did not specifically investigate this, our IHC data show high amyloid and tau accumulation in the EC and other regions of hippocampal formation. This however doesn’t preclude the validity of electrophysiology results since the data was collected specifically in the LEC region and it is unlikely that the neuronal dysfunction we see in the LEC is due to pathology in other brain regions.

Future research could further explore the mechanisms underlying the observed neuronal dysfunction in the LEC region. Previous studies have shown that fan cells are partially involved in object cell dysfunction and these specific cell-type vulnerability in AD could be studied. [49]. While this paper did not explore how LEC neuronal coding for odors is affected, other studies have indicated that fan cells are involved in associative memory and future studies could study if in AD these cells are affected [7]. In addition, given that the LEC is involved in episodic memories ([10]), it would also be interesting to investigate they affected by AD pathology.

Finally, the development and validation of LEC-specific memory tasks in human subjects could be a valuable avenue for early detection and intervention in AD.

## Materials and Methods

### Animals

We used a previously described transgenic AD mouse model, EC-APP/Tau ([3, 50]), and its littermates containing neuropsin promoter without amyloid beta and tau (NON) as age-matched control. These animals were used at 20+ months of age for behavior, 22+ months for electrophysiology experiments and euthanized before 25 months of age for immunohistochemistry. We used C57BL/6J mice (B6) as a young control for electrophysiology experiments. All animals were maintained on a 12-hour light/dark cycle with food and water provided ad libitum except for odor discrimination task where they were food restricted (details below). All animal experiments were performed in accordance with national guidelines (NIH; National Institutes of Health) and approved by the Institutional Animal Care and Use Committee at Columbia University.

### Behavior

#### Object Context Recognition

The experiment was performed during the light phase of the circadian cycle at a 30-lux light level in a 40-centimeter diameter cyilnder with interchangeable walls. Throughout testing, the experimenter remained unaware of the animal genotypes. Mice were tested in 2 different contexts; each displayed a pair of objects in a similar setup previously used for rats ([51]).

During training (sample phase), context 1 displayed white walls and presented a pair of small identical objects (objects 1A and 1B) whereas context 2 displayed gray walls and presented another pair of small identical objects (objects 2A and 2B). On habituation day (Day 1), all mice were allowed to explore both contexts without objects for 20 min in the morning and in the afternoon. During the first two days of training (Days 2 and 3), each animal was familiarized with both contexts and their pairs of identical objects. Mice were run 4 consecutive times for 3 min each time with rest intervals of 2 min in a holding cage. As a result, animals randomly experienced each context and their respective, identical objects twice per day. On testing day (Day 4), mice were randomly run in each context once before being tested in the last context to which they were exposed, only this time object 1A/B was substituted with object 2A/B or vice versa. The object that did not belong to the context was defined as the “novel object”, while the other object was referred to as the “familiar object”. Testing lasted 10 min and aimed to determine whether the animals spent more time exploring the novel object. A contingency table for entries into familiar object and novel object zone was tabulated and Fisher’s exact test was used to calculate the statistical significance. Odds ratio was calculated using Baptista-Pike method. Number of entries were converted to % entries and plotted as stacked bar graphs.

#### Odor Discrimination Task

The task was carried out with the same mice habituated in the same environment described in the T-maze spontaneous alternation protocol. Prior to testing, mice were single housed and food restricted. Weights were monitored and recorded in a logbook twice per day, before and after animal performance, to make sure mice remained above 85% of their body weight. Food restriction was adjusted accordingly. Animals were trained to discriminate between 2 distinct odors; odor A (Isoamyl acetate) and odor B (Ethyl valerate). Mice had to learn the association of a specific odor to a goal arm. To accomplish the task, animals performed a repetition of 40 trials per day for 4 consecutive days as follows: Each mouse was first confined in the stem where one of the 2 odors was presented to them using a Q-tip. Mice remained in the stem for 30 seconds, after which, the door was opened and the mice were allowed to choose an arm. On correct choices (for example, left arm for Isoamyl acetate), a reward (sugar pellet) was placed at the far end of the goal arm. After the animal consumed the reward, they were placed back in the stem of the T-maze. The maze was wiped with 70% ethanol in between each trial. The number of correct and incorrect entries into the arms for the final day were recorded in a contingency table. Data was analyzed using Fisher’s exact test to assess statistical significance.

#### T-maze Spontaneous Alternation

The experiment was conducted during the light phase of the circadian cycle at a 30-lux light level. Mice were tested to perform spontaneous alternation in a T-shaped maze as described previously ([52]). The maze measured 40 (stem) x 46 (arm) x 10 (width) cm and displayed a black cue card on the left side of the stem. During testing, experimenter was blind to the animal genotypes. On habituation day, all mice were allowed to explore both arms freely. Mice were then tested 6 times per day for 3-4 consecutive days as follows: Each mouse was first placed at the stem and offered to explore one randomly-picked open goal arm (with other arm closed) for 1 min. If the goal arm was reached, mice were placed at the stem again and the trial was repeated with both arms open. Testing was ended after the animal chose one arm or stayed in the stem for 1 min. Between trials, the maze was wiped with 70% ethanol. The number of correct and incorrect choices for all days (alternative goal-arm and same-arm/no-arm choices) were recorded in a contingency table. Data was analyzed using Fisher’s exact test to assess statistical significance.

#### Surgery and electrode implantation

Custom-made, reusable 16-channel microdrives (Axona, UK) were constructed by attaching an inner (23 ga) and an outer (19 ga) stainless steel cannula to the microdrives ([53]). Tetrodes were built by twisting four 25 mm thick platinum-iridium wires (California wires) and heat bonding them. Four such tetrodes were inserted into the inner cannula of the microdrive and connected to the wires of the microdrive. One day prior to surgery, the tetrodes were cut to an appropriate length and plated with a platinum/gold solution until the impedance dropped to about 150 kohms. On the day of surgery, mice were anesthetized with a mixture of ketamine and xylazine (100 mg/ml and 15 mg/ml, respectively, per 10 g body weight) and monitored for depth of anesthesia before proceeding. Mice were then fixed within the stereotaxic frame with the use of zygomatic process cuff holders and an incision was made to expose the skull. About 3–4 jeweler’s screws were inserted into the skull to support the microdrive implant. An additional screw connected with wire was also inserted into the skull which served as a ground/reference for local field potential (LFP) recordings. A 2 mm hole was made on the left hemisphere of the skull at ML 4.3 mm and AP -3.4 mm from bregma and tetrodes were lowered to -3.0 mm to target the LEC. Dental cement was spread onto the skull and secured with the microdrive. Mice were allowed to recover from anesthesia in a cleaned cage placed on a warm heating pad until awake (45 min) before finally being transported to housing. Carprofen (5 mg/kg) was administered to mice prior to surgery and post-operatively to reduce pain. Mice usually recovered within 24 h, after which the tetrodes were lowered and recording began. All recording depths reported are from the surface of the brain.

#### In vivo electrophysiology recording and cell classification

The mice explored a 40 cm diameter white cylindrical arena with a vertical black cue card (8.5 × 11 inch) attached to north-side of the cylinder. Mice underwent 3-5 recording sessions per day, with 10 minutes to 2 hours between sessions. A session always started without an object in the cylinder (No object) followed by an object positioned at the edge of the cylinder at either North (0 degrees), East (90 degrees), South (180 degrees) or West (270 degrees) direction. Tetrode positions were moved no more than 50*µ*m at a time, and only after the last recording session of the day, allowing 12 hr of stable electrode positioning prior to the next recording session.

Neuronal signals from experimental mice were recorded using the Axona DacqUSB system. The signals were amplified 15,000 to 30,000 times and band pass filtered between 0.8 and 6.7 kHz. LFP was recorded from 4 channels of the electrodes. The LFP was amplified 15,000 times, lowpass filtered at 125 Hz and sampled at 250 Hz. Notch filter was used to eliminate 60 Hz noise. The recording system tracked the position of the infrared LED on the head stage (sampling rate 50 Hz) by means of an overhead video camera. Tracking artifacts were removed by deleting samples greater than 100 cm/s and missing positions were interpolated with total durations less than 1 s, and then smoothing the path with a 21-sample boxcar window filter (400 ms; 10 samples on each side). Spike sorting was performed offline using MountainSort software ([54]) followed by manually checking and validating the clusters. Quantitative measurements of cluster quality were subsequently performed, yielding isolation Distances in Mahalanobis space ([55]). There were no significant differences between clusters in our experimental groups (Median Isolation distances: 8+ mo B6-9.44, 24+ mo NON-10.13 and 24-mo EC-App/Tau-9.83. Kruskal-Wallis test-P = 0.334). A total of 704 neurons from 15 animals were recorded with 199 neurons in B6 mice (N=5), 169 in NON mice (N=5) and 336 in EC-App/Tau mice (N=5). To match cells across sessions, a custom-script in Python called Unit matcher was used which compared the waveform properties of all cells and re-assigned the cell numbers in the clusters.

#### Neural coding properties

All the data for neural coding properties (e.g. information content) are expressed as the mean ± the standard error of the mean (SEM). We performed the D’Agostino & Pearson omnibus normality test to determine if the data was normally distributed. We then chose the following statistical tests. Kruskal-Wallis test was used to compare medians in all groups. Fisher’s exact test was used to compare the binary data from the Object context recognition, Odor discrimination and T-maze tests. A value of *p* < 0.05 was considered statistically significant for all measures. The N represents the number of animals in each group, unless specified otherwise. The exact values of N are indicated in figure legends, and in the Methods section.

#### Object and trace cell analysis

We used our previously published EMD method [42] to calculate object and trace cell stability across sessions. For a given cell’s ratemap on a given session, we quantified remapping relative to chosen reference points. Specifically, we computed a remapping distance between an activity map and the object location. We did this for the full ratemap, to capture whole map stability, and for a masked version, to isolate and track individual fields. We used the single point Wasserstein distance (normalized EMD distance), a strong approximation of the true distance between the firing rate map and a full pseudo-rate map with all the density at one point.

Alongside these EMD metrics, we also compute a euclidean distance between the weighted average firing peak of a given field and each of the reference points. We refer to this as the centroid distance.

#### Reference points and distribution

The reference points were selected as the 4 tested object locations (0, 90, 180, 270). We also computed a 5th reference point set at the center of the arena (NO = no object). While an object may or may not have been present on a given session, we computed remapping relative to all 5 reference points for every cell ratemap on every session. This allows us to track object/trace behavior and movement towards the objects throughout tested sessions.

For each reference point we compute a remapping distance and convert this into a quantile based on a reference distribution. For a given cell on a given session, we computed the minimum and maximum ‘x’ and ‘y’ positions covered by the animal on that session. This gave us an egocentric map of possible (x,y) locations covered. Given the EMD property where gradients of remapping emerge across the map when localizing a field, we decided to sample in a way that would give us coverage across these gradients. We therefore sampled from the possible (x,y) positions using a hexagonal lattice with a spacing parameter of 2 resulting in a hexagon side of 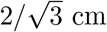. Points were sampled from all 6 vertices of each hexagon as well as the center. We also masked the collected points to ensure only those that would fall inside the circular arena were selected. This gave us a uniform sampling across the gradients and on average consisted of 776 ± 148 sample points for a given map.

Using these sample points, we computed a remapping distance from the given map score to each of these randomly sampled reference points. With a distribution of remapping distances to random reference points and our empirically observed distances (for each of the 5 reference points), we compute a quantile describing the spatial location/dispersion of our map/field relative to other possible random locations.

#### Cell assignments

We define cells in three categories: object, trace and unassigned. Each cell appearance in a session is labeled with isTrace and isObject identifiers, which are set to 1 if conditions are met, otherwise 0. For object cells, if the minimum quantile during a session matches the current location, isObject is set to 1. For trace cells, isTrace is 1 if the quantile matches any prior tested location. After processing all cell appearances, we apply filters to ensure data integrity and exclude ambiguous cells. We use three filters:

- Exclude cells with an isolation distance under 5.
- The quantile using the arena center must exceed the quantile of at least 1 object angle. In other words, the arena center quantile cannot be the minimum
- The reference point of the minimum quantile using the field-restricted EMD should align with the minimum quantile using the centroid distance. This accounts for cases where two object angles have very similar quantiles.

The first filter is applied across all cell appearances. The next two filters are used to generate filtered cell assignments based on the ambiguity of a field’s position. These two filters are relative in that they require no threshold. We preserve cells appearing in at least two sessions post-filtering. Definitions applied for cell assignment are:

- An object cell appears in at least two sessions with the minimum quantile at the current location.
- A trace cell appears in at least one session with the quantile at a prior object location.

Object cells are prioritized over trace cells, minimizing their overlap (1-2 cells typically). This methodological rigor helps maintain the session structure and prevents loss of data from low-quality recordings.

We performed Fisher’s exact test to determine if there were differences between the total population cell assignments (object, trace, unassigned). We then performed pairwise 2×2 Fisher’s exact test to compare specific group pairs within a given assignment of object or trace. The 2×2 Fisher’s exact tests were also applied separately in each group to group comparison for each of the quality metrics.

#### Spatial stability

To explore spatial stability over time (relative to the arena center), we grouped our data by session number from 1 to 6 and computed a mean quantile for each group. There were 6 means for ANT and NON and 3 means for B6. The Mann-Kendall test was applied to obtain an empirical test statistic for each. The slope was computed as the Theil-Sen estimator. To assess significance given the low number of sessions and high fluctuations within a given session we applied a bootstrap Mann Kendall test where, for a given session number, we shuffled data points across the 3 groups and recomputed a new set of mean values from which a Mann Kendall test statistic was computed. This was done 1000 times to create a distribution to which the empirical test statistic was compared and a p-value obtained. Benjamini-Hochberg corrections were applied to collected p-values.

To look at overall group stability, we used the standardized EMD quantiles as part of a beta regression model to compare quantiles across two groups. For each possible group to group comparison, we fit a model with the group id and session id as fixed effects, the animal id as a random effect and a session id to group id interaction term.

This gives the formulas:

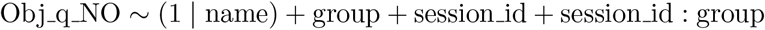

#### DAB Immunohistochemistry and Imaging

After final recording, all mice were anesthetized with a mix of ketamine/xylazine and were transcardially perfused with ice-cold 100 mM phosphate-buffered saline (PBS, pH 7.4), followed by 10% formalin (Fisher Scientific, USA). For mice implanted with microdrive, the electrodes were not moved until after perfusion. Brains were then harvested and left in 10% formalin overnight, then incubated in 30% sucrose until the brains sank to the bottom of the conical tube (all at 4°C). Horizontal brain sections were sliced (30 *µ*m) using a Leica CM3050 S cryostat and stored in cryoprotectant at -20°C until immunostaining procedures. The EC-APP/Tau mice brain sections were processed in parallel with sections from NON and B6 mice where appropriate.

Immunoperoxidase staining was performed using a Mouse-on-Mouse kit (Vector Laboratories, USA) as described previously [46]. Briefly, free-floating tissue sections were quenched with 3% H2O2 and blocked with mouse IgG–blocking reagent for 1.5 hours at room temperature, followed by overnight incubation at 4°C with either anti-beta amyloid 6E10 (mouse, 1:1,000 dilution of 1 mg/mL stock) (Biolegend, USA), anti-tau MC1 (mouse, 1:500; courtesy of Peter Davies, Feinstein Institute for Medical Research, North Shore LIJ), biotinylated anti-phospho tauSer202/Thr205 AT8 (mouse, 1:500) (Thermo Scientific, USA) or anti-human tau CP27 (mouse, 1:500; courtesy of Peter Davies). Tissue sections were then rinsed in PBS and incubated for approximately 20 minutes at room temperature in a working solution of biotinylated anti-mouse IgG reagent (excluding the biotinylated AT8-labeled sections). After several PBS rinses, sections were then incubated in an avidin-biotin conjugate for 10 minutes before being developed in H2O containing 3,3’-diaminobenzidine (DAB) hydrochloride and urea hydrogen peroxide (Sigma Aldrich, USA). After staining was completed, tissue sections were mounted onto glass Superfrost Plus slides (Fisher Scientific, USA), allowed to air dry completely, and then dehydrated in ethanol and cleared with xylenes before being coverslipped. Tissue sections were analyzed under bright-field microscopy using an Olympus BX53 upright microscope. Digital images were acquired using an Olympus DP72 12.5 Megapixel cooled digital color camera connected to a Dell computer running the Olympus cellSens software platform (Olympus Corporation; https://www.olympus-lifescience.com/en/software/cellsens/).

#### Code

Custom Python codes were used to match units across sessions, quantify neural coding, assess spatial stability and identify cell types (NeuroSciKit). Post-hoc analyses and visualizations were done on Jupyter notebooks and using Matplotlib or Prism 9 software (GraphPad, San Diego, CA, USA).

## Author contributions

S.A.H. performed the surgeries and implanted the electrodes. M.H., E.N. and S.A.H. collected the behavioral and electrophysiology data. M.H. and E.N. performed immunohistochemistry.

R.R. and S.A.H. sorted the spike data. R.R. and S.A.H. analysed the behavioral and immunohistochemistry data. A.A. and O.S. developed the analytical code in Python. A.A. and R.R. processed the electrophysiology data. A.A. conducted post-hoc analyses, with help from O.S., and data visualizations. S.A.H. conceived the idea, discussed the methods and results with R.R, A.A. and M.H.. R.R., A.A., M.H. and S.A.H. wrote the manuscript. S.A.H. supervised the project and obtained the funding.

## Acknowledgements

We would like to thank Dr. Karen Duff for providing us with the EC-App/Tau line along with neuropsin controls. We thank Aisha Kazeem, Lisa Shah, Adithi Jayaraman and Sandhya Senthilkumar for help with spike sorting earlier dataset that first identified object and trace cells. We thank Dr. Gustavo Rodriguez for help with immunohistochemistry techniques provided to M.H. and for the valuable feedback on the manuscript. These tools were supported by NIH/NIA grants R01AG050425 and R01AG064066.

## Supplementary Figures

**Supplementary Figure 1.**
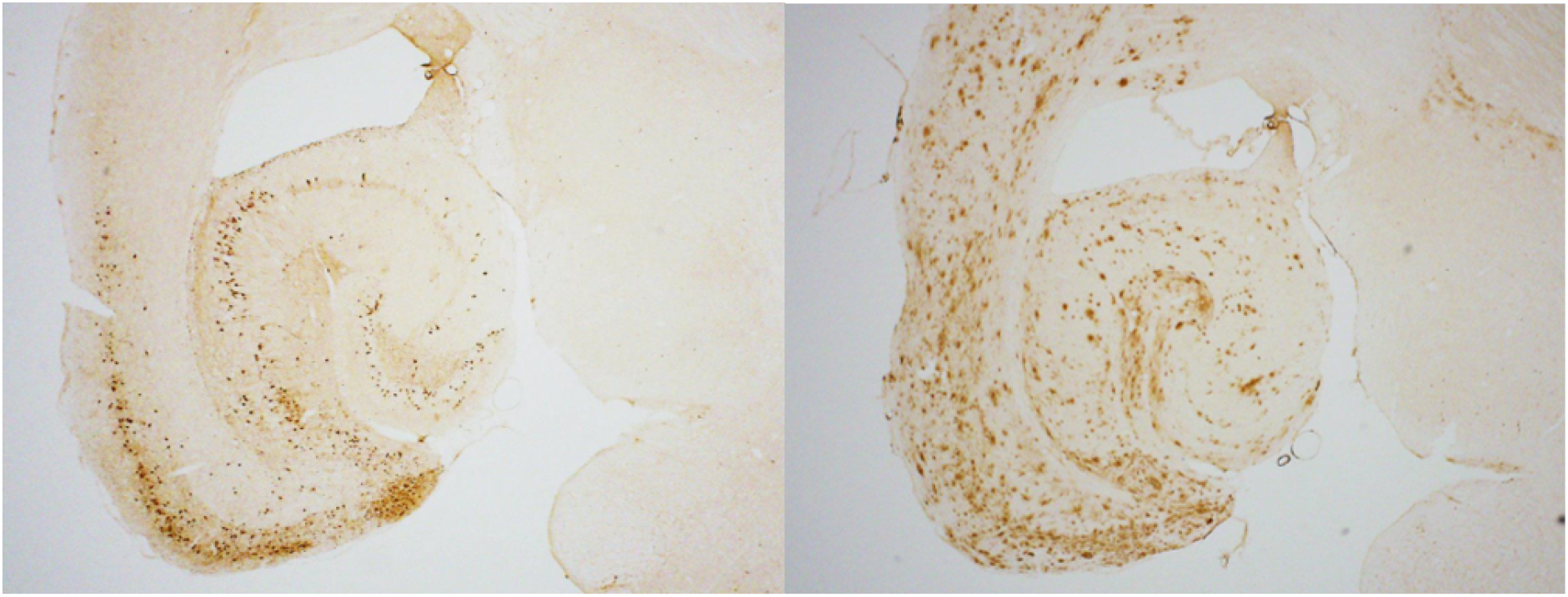
Tau and Amyloid beta accumulation in EC-App/Tau mice. **Left**: a representative horizontal tissue section from a 24-month-old EC-App/Tau mouse with pronounced tau accumulation (AT8 staining) in the EC region and some in the neighboring hippocampal regions. **Right**: A*β* accumulation in the hippocampal formation of same mouse as left with more deposition in the EC region

**Supplementary Figure 2.**
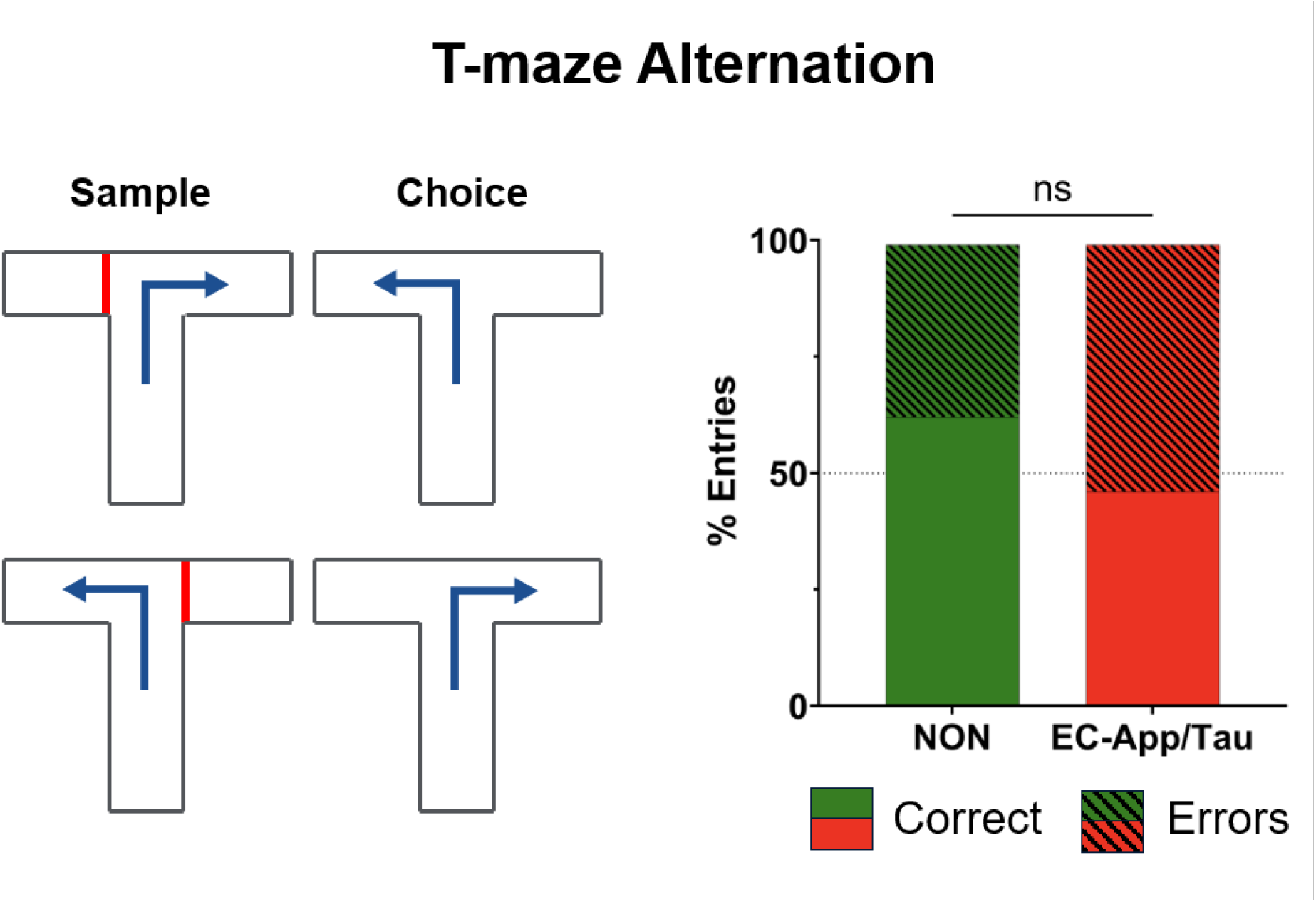
EC-app/Tau mice are not impaired in T-maze alternation task. **Left**: Schematic of two trial blocks with sample (left) and choice (right) runs. **Right**: Stacked bars showing relative % entries for NON and EC-App/Tau mice into left or right arm of the T-maze. EC-App/Tau mice made fewer correct entries and more errors compared to the NON mice but this was not significant.

**Supplementary Figure 3.**
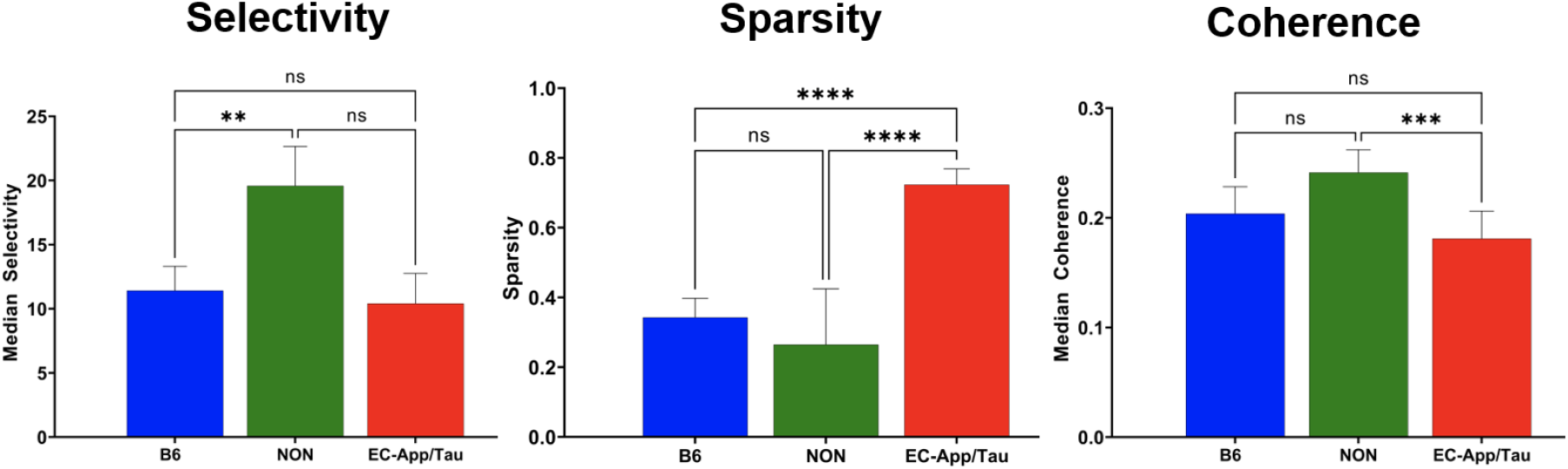
Selectivity, Sparsity and Coherence The histogram on the left represents comparisons amongst the B6, NON and EC-App/Tau cohorts for selectivity; the panel in the middle shows sparsity and that on the right shows coherence. statistical significance is represented by asterisks wherein ns indicates P*>*0.05, ** indicates P<0.01, *** indicates P< 0.001 and **** indicates P<0.0001

**Supplementary Figure 4.**
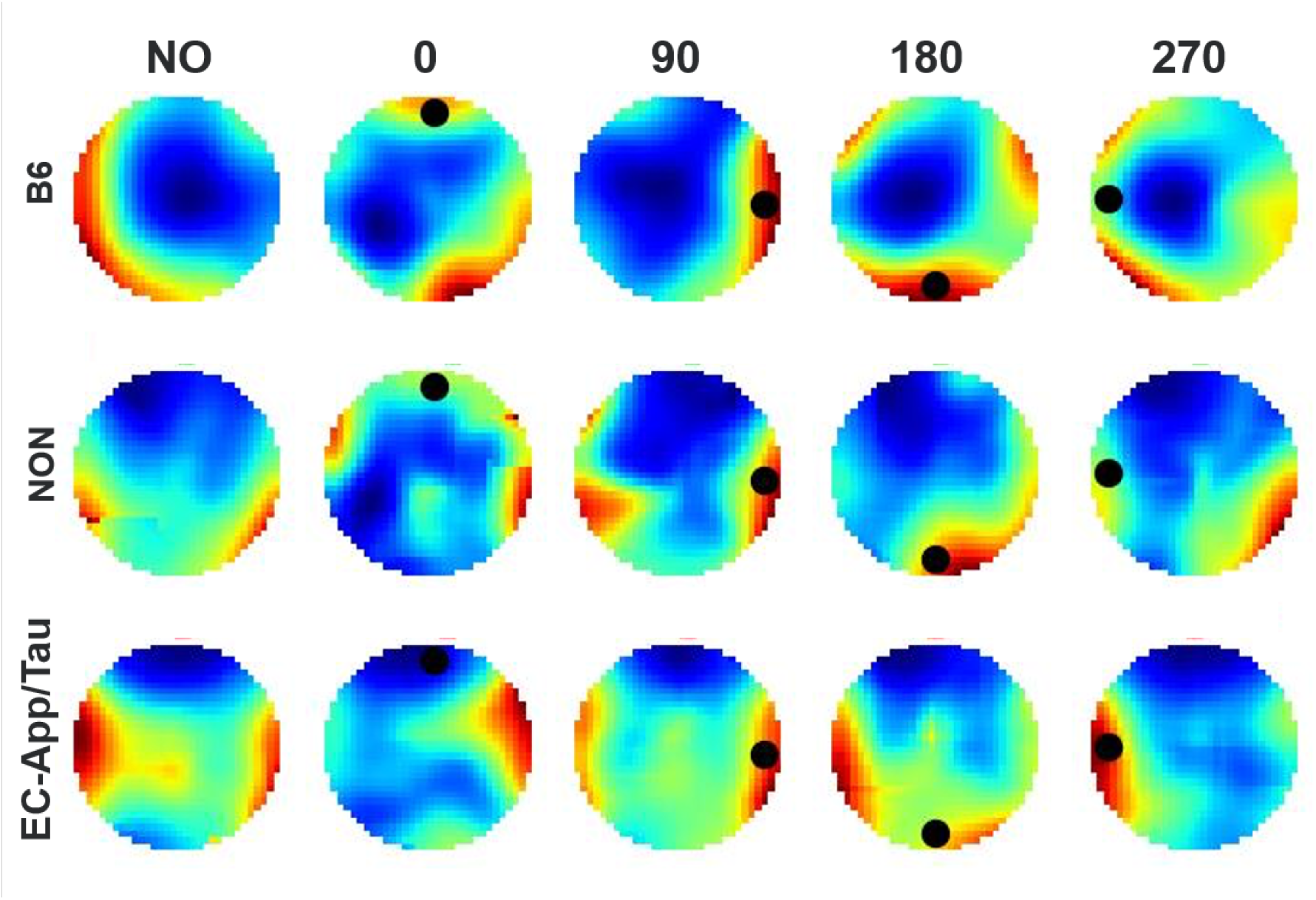
Average population responses. All valid cells are considered across 3 groups (B6, NON, EC-App/Tau) and 5 possible locations (0, 90, 180, 270, NO = no object). Ratemaps averaged by mouse group (‘B6’, ‘NON’, ‘EC-App/Tau’) (A). Ratemaps grouped by current object location on a given session and averaged (B). The top row displays a representative sequence with current (black) and previous (white) object locations displayed at each step.

**Supplementary Figure 5.**
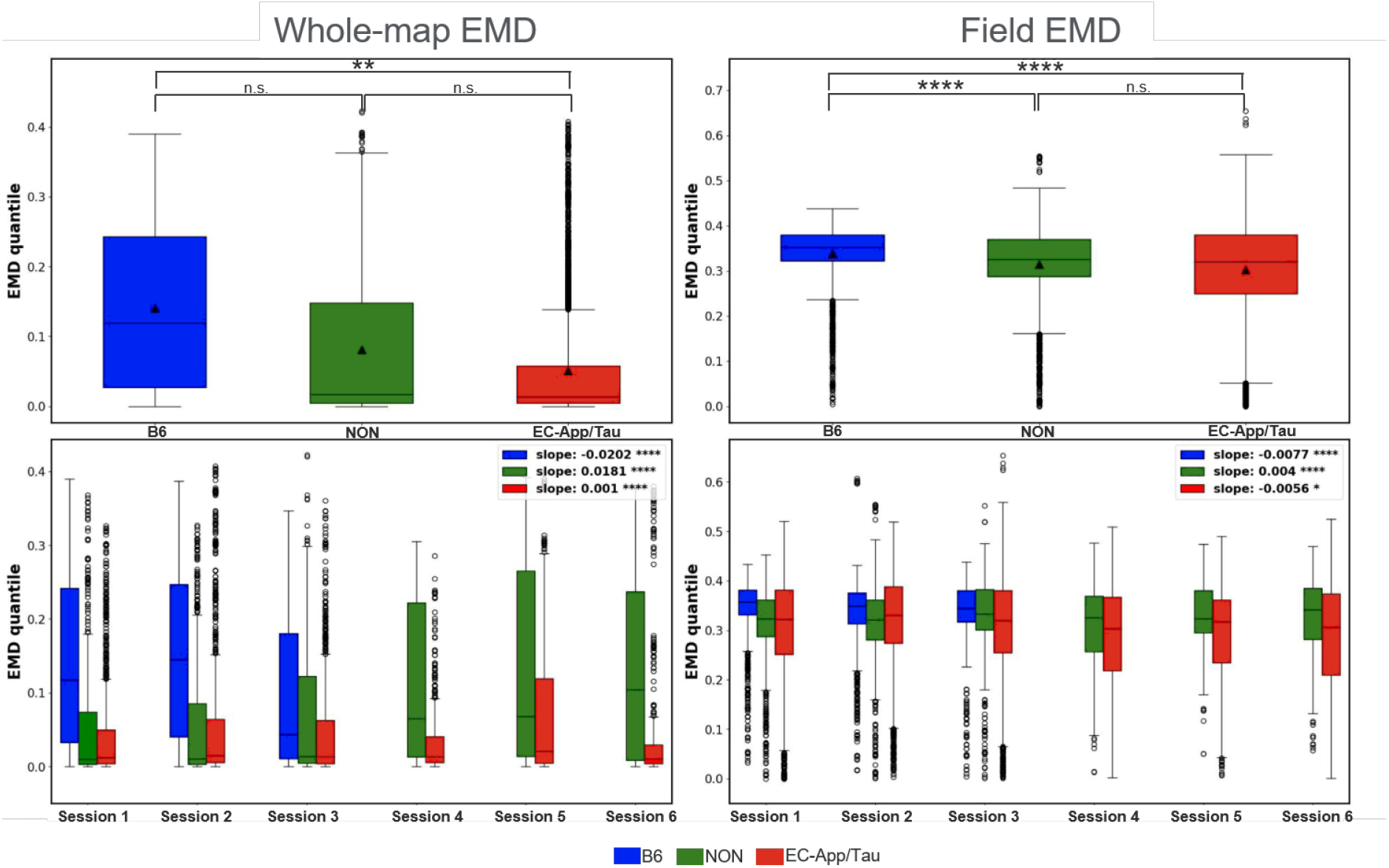
Group stability. Stability metrics are displayed as quantiles for the whole map EMD (left column) and the field-restricted EMD (right column). The 3 groups (B6, NON, ANT) are compared using boxplots with significance values from the regression group coefficient (top row). EMD quantiles are separated by session number and trends are compared using a bootstrapped Mann-Kendall slope test (bottom row).

**Supplementary Figure 6.**
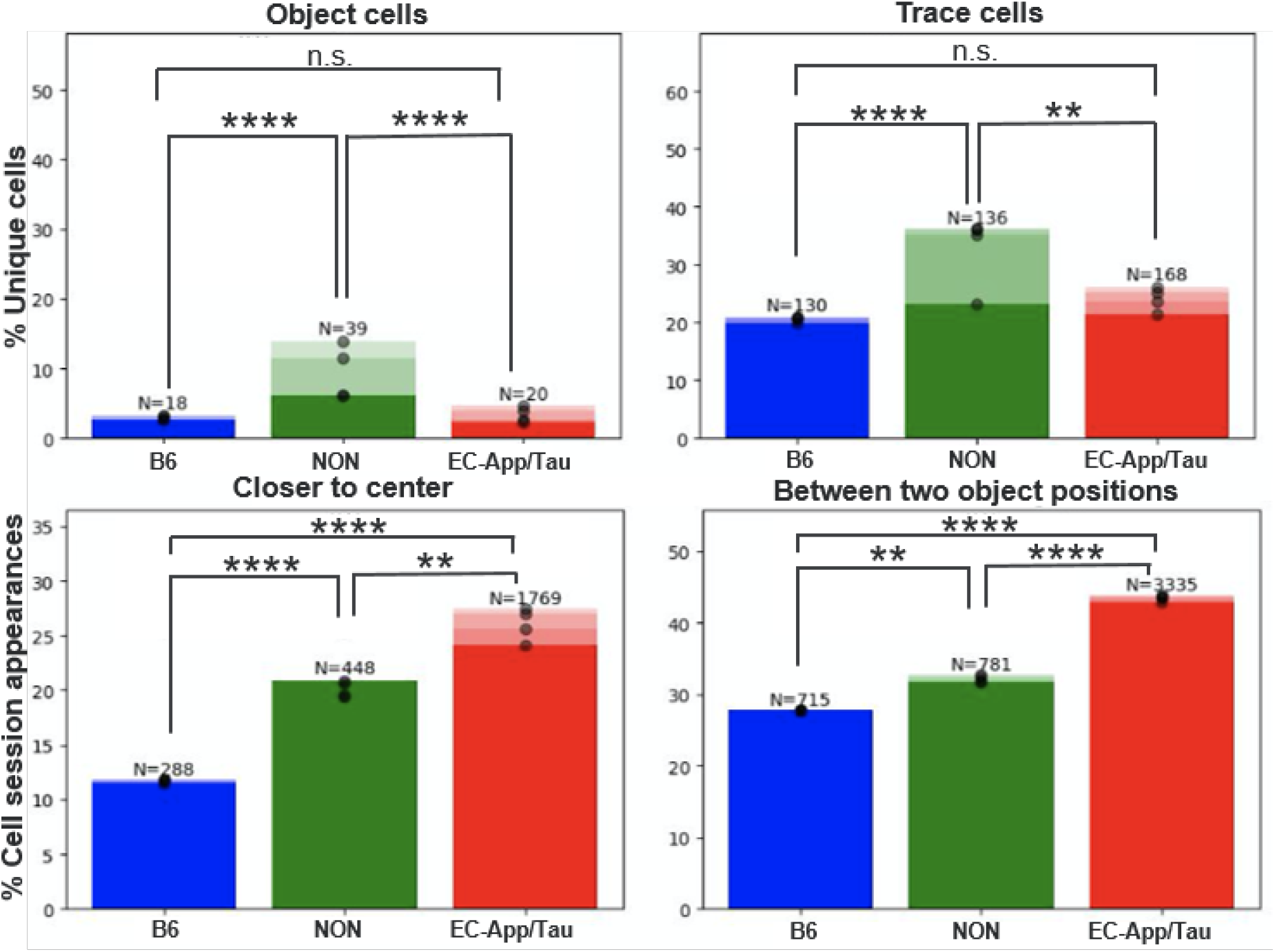
Cell Counts Over Time. Cell counts are compared using Fisher’s exact test. The percentage of filtered cells which show object and trace behavior with an increasing session limit compared for the 3 groups (Top panel). Later sessions are shaded in progressively lighter colors. The percentage of cell-session appearances (individual ratemaps) failing a quality metric are displayed (Bottom panel). The first quality metric shows the proportion of cases where the EMD quantile is smaller relative to the center point than to any of the 4 object locations (Bottom left). The second shows the proportion of cases between two object locations such that the reference point of the minimum quantile is different for the field-restricted EMD and the field centroid distance (Bottom right). Raw cell and cell-session counts are provided above each proportion.

**Supplementary Figure 7.**
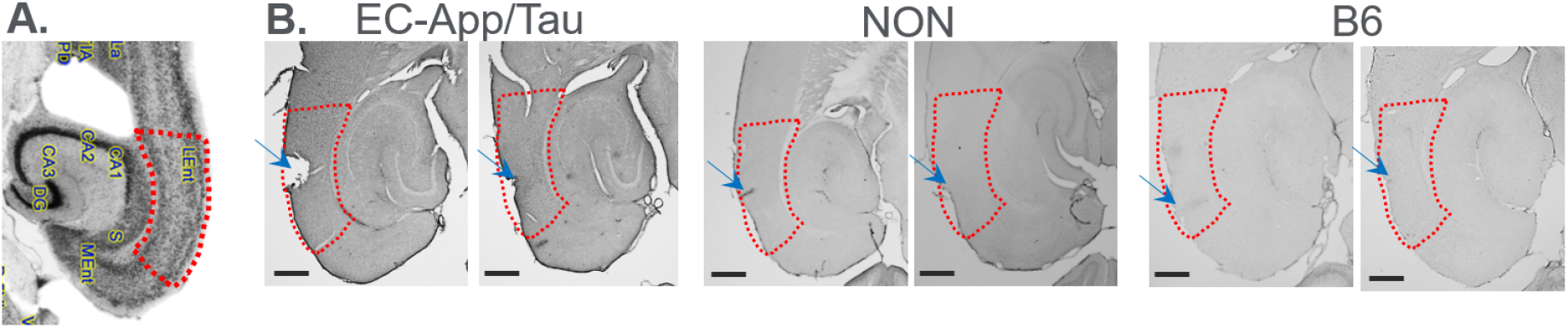
Tetrode position at LEC. **A**. Horizontal section of B6 mouse taken from Mouse Brain Library (Rosen et al., 2000) showing different sub-regions within the hippocampal formation including the LEC (labelled as LEnt within the red inset). **B**. Representative sections from each group; EC-App/Tau, NON and B6 mice, showing tetrode holes/marks (blue arrows) within the LEC region (red inset). Two animals per group were stained with AT8 antibody to detect phosphorylated tau. As expected, left two sections (EC-App/Tau mice) show tau deposits in the LEC while NON and B6 show no tau deposition. Black scale bars at left bottom are 500 *µ*m.

